# Mechanical activation of epithelial Na^+^ channel relies on an interdependent activity of the extracellular matrix and extracellular *N*-glycans of αENaC

**DOI:** 10.1101/102756

**Authors:** Fenja Knoepp, Zoe Ashley, Daniel Barth, Marina Kazantseva, Pawel P. Szczesniak, Wolfgang G. Clauss, Mike Althaus, Diego Alvarez de la Rosa, Martin Fronius

## Abstract

Mechanotransduction describes how cells perceive their mechanical environment and mechanosensitive ion channels are important for this process. ENaC (epithelial Na^+^ channel)/DEG (degenerin) proteins form mechanosensitive ion channels and it is hypothesized their interaction with the extracellular matrix (ECM) *via* ‘tethers’ is required for mechanotransduction. Channels formed by vertebrate α, β and γ ENaC proteins are activated by shear force (SF) and mediate electrolyte/fluid-homeostasis and blood pressure regulation. Here, we report an interdependent activity of ENaC and the ECM that mediates SF effects in murine arteries and heterologously expressed channels. Furthermore, replacement of conserved extracellular *N*-glycosylated asparagines of αENaC decreased the SF response indicating that the attached *N*-glycans provide a connection to the ECM. Insertion of *N*-glycosylation sites into a channel subunit, innately lacking these motifs, increased its SF response. These experiments confirm an interdependent channel/ECM activity of mechanosensitive ENaC channel and highlight the role of channel *N*-glycans as new constituents for the translation of mechanical force into cellular signals.

## Introduction

Mechanotransduction describes how cells perceive their mechanical environment by translating mechanical forces into cellular signals. Among other transmembrane proteins, mechanosensitive ion channels are key molecules for this process (Hamill & Martinac, 2001; Jaalouk & Lammerding, 2009). Currently, two distinct principles have been put forward to explain how force can activate mechanosensitive ion channels: 1) the ‘force-from lipids’ (Teng et al, 2015) and 2) the ‘force-from filaments’ principles (Katta et al, 2015). Both principles are based on the assumption that mechanical force is sensed and transmitted by a specialized ‘sensing structure’ to induce a conformational change in an ion channel as the ‘receiver molecule’ (Suppl Fig 1). The force-from filaments principle (also known as the ‘tethered model’), proposes the transmission of mechanical force through tethering of the ion channel proteins to internal (cytoskeletal filaments) and/or external filaments (extracellular matrix: ECM) as ‘sensing structures’ (Gillespie & Walker, 2001; Zanini & Göpfert, 2013; Krieg et al, 2015). Deformation or deflection of the sensing structure will be transduced *via* tethers to the channel to change its conformation and thereby its activity.

**Figure 1.**
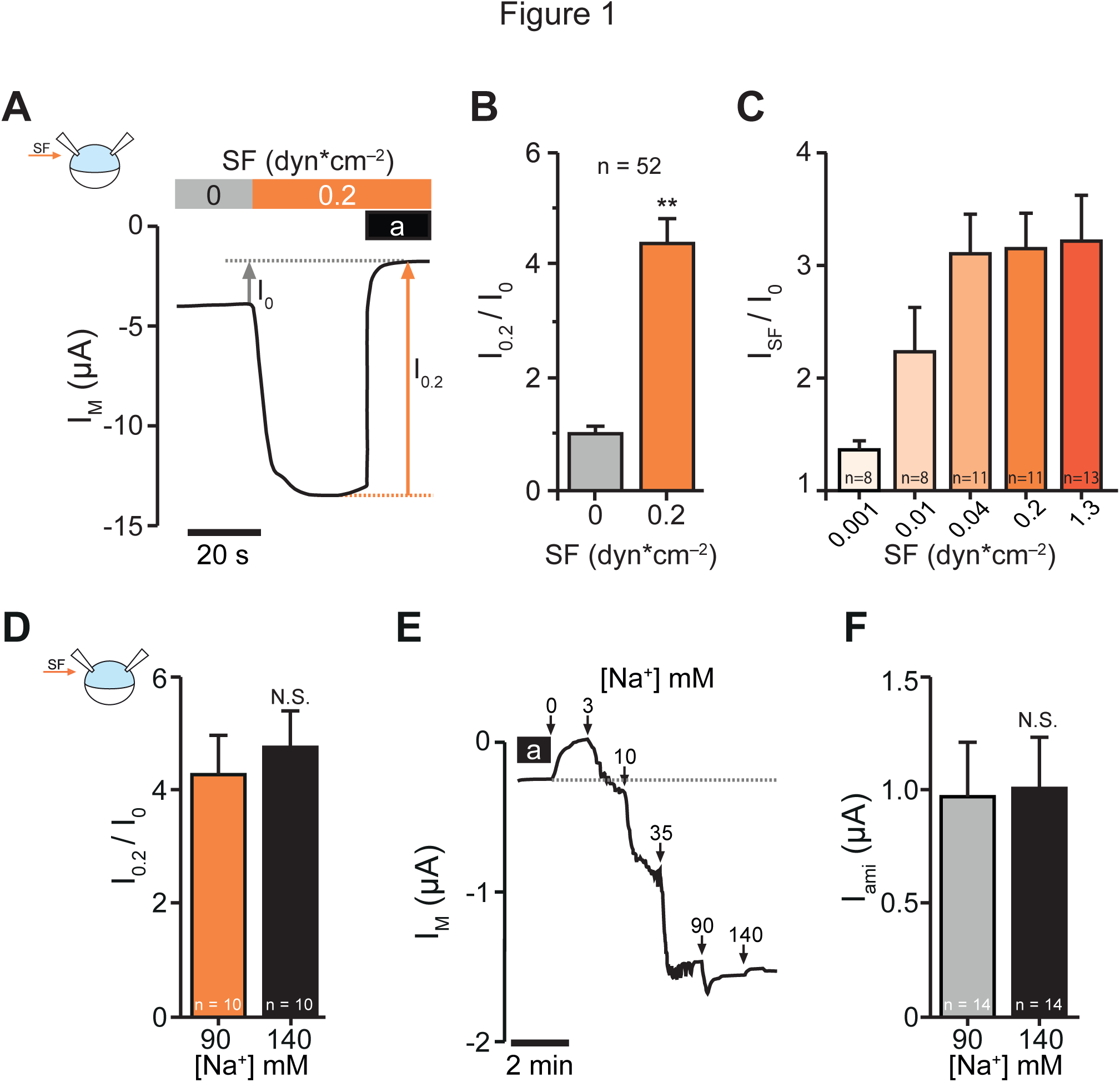
**ENaC activation by SF is independent of an unstirred layer effect.** A Application of SF by turning on the bath perfusion activates αβγENaC. Without SF (0 dyn*cm^−2^, grey bar) a certain baseline ENaC activity was observed. Activating the bath perfusion (0.2 dyn*cm^−2^, orange bar) induced a rapid increase of the transmembrane current (I_M_). Amiloride (a, black bar) was applied to estimate ENaC-mediated current in the absence (I_0_) and presence of SF (I_0.2_). B Current values with SF (I_0.2_) were normalized with respect to the values before SF was applied (I_0_). (**: *P* < 0.01, one sample *t*-test, two tailed). C The SF-response (I_SF_) was augmented by increases in SF. D SF-responses (I_0.2_/I_0_) were not influenced by extracellular Na^+^-concentrations (90 and 140 mM). Osmolarity was adjusted with mannitol (N. S., not significant, two tailed Mann-Whitney test). E Dose response curve for extracellular Na^+^, performed on an αβγENaC-expressing oocyte which was continuously exposed to low SF (0.01 dyn/cm^2^). The incremental enhancement of extracellular Na^+^-concentrations from 0 to 90 mM induced a stepwise increase of I_M_, but the elevation from 90 mM to 140 mM Na^+^ had no further effect. The dashed line indicates the amiloride sensitive baseline current (I_ami_). Differences in osmolarity between the solutions were compensated by appropriate amounts of mannitol. F ENaC-mediated currents with 90 mM and 140 mM extracellular Na^+^ are similar. N. S., not significant, two tailed unpaired *t*-test.

There is emerging evidence for intracellular and extracellular tethers regulating mechanical activation of channels. For example mechanical activation of No Mechanoreceptor Potential C (NOMPC) channel of *Drosophila melanogaster* depends on ankyrin repeats within its *N*-terminus that form an intracellular ‘tether’ (Zhang et al, 2015), likely to facilitate the connection to microtubules as identified for the transient receptor potential vanilloid type-1 (TRPV1) channel (Prager-Khoutorsky et al, 2014) that belongs to the same protein family as NOMPC. Evidence for extracellular tethers as constituents of mechanotransduction originates from studies in *Caenorhabditis elegans* and hair cells (Gillespie & Walker, 2001; Chalfie, 2009). In *C. elegans* mutations of genes encoding ion channel subunits (e.g. mechanosensory abnormality-4 (MEC-4) and MEC-10) and extracellular proteins (e.g. MEC-5 and MEC-9, that are considered to be part of the ECM) resulted in touch-insensitive animals (Du et al, 1996; Emtage et al, 2004). This indicates that channel proteins and the ECM are connected – *via* tethers – to facilitate the mechanical perception of touch. In hair cells extracellular tethers known as tip links have been identified by electron microscopy (Pickles et al, 1984; Furness & Hackney, 1985) and there is evidence that such filamentous protein structures are directly involved in channel gating of neurons that mediate touch sensation (Hu et al, 2010).

The mechanosensitive MEC and DEG channel proteins of *C. elegans* are the founding members of the epithelial Na^+^ channel (ENaC)/degenerin (DEG) protein family. Members of this family are widely expressed in the animal kingdom and form mechanosensitive ion channels (Kellenberger & Schild, 2002; Hanukoglu & Hanukoglu, 2016). Well-known members of the family constitute the epithelial Na^+^ channel (ENaC) in vertebrates (Canessa et al, 1993; Chalfie et al, 1993). Canonical ENaC is composed of α-, β- and γ-subunits (Canessa et al, 1994b) (SCNN1A, SCNN1B and SCNN1G), highly abundant in absorbing epithelia and is crucial for electrolyte/fluid-homeostasis (Rossier et al, 2015) and blood pressure regulation (Warnock et al, 2014). Soon after discovering the molecular identity of ENaC, evidence emerged that ENaC is mechanosensitive and activated by membrane stretch (Awayda et al, 1995; Achard et al, 1996; Ismailov et al, 1997; Ji et al, 1998), in accordance with the ‘force from lipids’ principle. However, these studies were critically discussed (Rossier, 1998) suggesting that there is not enough evidence for the conclusion that ENaC is mechanically activated by stretch.

In contrast to the controversial findings about the activation by membrane stretch, there is compelling evidence that ENaC is mechanically regulated by shear force (SF) (Satlin et al, 2001; Carattino et al, 2004; 2005), that is defined as force acting parallel on a surface (Suppl Fig 1). An example for the generation of SF is represented by the blood flow, when passing the luminal surface of arteries. The SF-mediated effect on ENaC was confirmed in isolated kidney tubules (Morimoto et al, 2006), and in outside-out single channel recordings where an increased open probability was observed in response to increased SF of expressed (Althaus et al, 2007) and endogenous channels (Wang et al, 2009). The SF-mediated regulation of ENaC is of considerable importance for blood pressure regulation (Warnock et al, 2014). ENaC subunits expressed in endothelial cells of arteries (Golestaneh et al, 2001), arterial baroreceptors (Drummond et al, 1998; 2001) and distal kidney epithelium (Loffing et al, 2001) are permanently exposed to SF due to the fluids flowing over the surface of the cells. This indicates that ENaC’s ability to sense and transduce SF is vital for its function and that it is an ideal candidate to study the impact of SF on channel activity.

Since ENaC proteins are related to the mechanosensitive proteins of *C. elegans* it was hypothesized that mechanical activation involves the ECM as well as extracellular ‘tethers’ that facilitate the interdependent activity (Drummond et al, 2008; Fronius & Clauss, 2008) – in agreement with the ‘force-from filament’ principle. Although the ‘force-from filament’ principle is well accepted for ENaC/DEG channels, experimental evidence is scarce. In addition, little is known about the architecture of the hypothesized extracellular tethers. Therefore, our study focuses on two aspects: (1) Provide direct functional evidence for the interdependent activity of the channel and the ECM for mechanical activation. (2) Identifying potential interaction sites of channels that are required for the interaction with the ECM.

The role of the ECM for SF sensation was addressed by electrophysiological experiments in *Xenopus laevis* oocytes representing an expression system with an intact ECM and pressure myography on isolated perfused murine carotid arteries. In both systems, enzymatic ECM-degradation impaired the SF responses of ENaC. Further, we targeted the role of *N*-glycans attached to extracellular asparagines of the channel as potential components of extracellular tethers involved in facilitating the connection between the channel and the ECM. By removal of certain *N*-glycosylated asparagines the SF response was decreased, whereas the addition of *N*-glycosylation motifs increased SF responses. Thus we provide evidence that an intact ECM and channel *N*-glycans are new interdependent constituents for ion channel mechanotransduction.

## Results

### SF-dependent activation of ENaC involves the extracellular matrix

SF-dependent activation of expressed human αβγENaC was analyzed by performing electrophysiological recordings on *Xenopus laevis* oocytes (Fig 1A-C) and HEK293 cells (Suppl Fig 2). Although we observed SF-induced increases in amiloride-sensitive current in ENaC-expressing HEK293 cells, the SF-induced effects were inconsistent (Suppl Fig 2). Transmission electron microscopy combined with slam freezing fixation of HEK293 cells revealed a fibrous matrix on the luminal side of HEK cells (Suppl Fig 4), although this matrix varied between different passages. This finding indicates that the ECM of HEK293 cells undergoes changes during passaging for cultivation and that these changes may account for the inconsistent SF effects. Since a native and consistent ECM was crucial for this study *Xenopus* oocytes were used for ENaC expression. In contrast to other cell lines, oocytes have a native and differentiated ECM that is not challenged by passaging during cultivation. They are also an established model for investigating the effects of SF on ENaC (Carattino et al, 2004; Althaus et al, 2007), K^+^ channels (Hoger et al, 2002), and MEC channels (Shi et al, 2016).

In *Xenopus* oocytes – as previously reported – an ENaC-dependent increase of the transmembrane current (I_M_) was observed by the application of SF (Fig 1A-C). A SF response was not observed in the presence of amiloride (10 µM) or in water-injected oocytes (Suppl Fig 3). Further experiments were performed to assess whether an unstirred layer effect influences the SF response of ENaC due to the different kinetics of Na^+^ supply to the channel surface (Mackay & Meares, 1959). Therefore, SF responses with 90 and 140 mM extracellular Na^+^ were recorded and compared. The observed SF responses were similar (Fig 1D) and almost identical compared with the effects shown in Fig 1A and B. Additionally, dose response curves with increasing extracellular Na^+^ were performed to observe if 140 mM Na^+^ alters ENaC currents in general. As shown in Fig 1E and F 140 mM Na^+^ had no additional effect. These experiments confirm that the SF-effect is not affected by an enhanced Na^+^ supply and that an unstirred layer effect can be excluded.

To determine a direct contribution of the ECM for SF-sensing of expressed ENaC, the channel’s response to SF was assessed upon degradation of oocyte ECM-components. In oocytes, the ECM consists of 1) the vitelline envelope representing the outer surface of defolliculated oocytes, and 2) a filamentous glycocalyx within the perivitelline space which is between the vitelline envelope and the oocyte’s membrane (Larabell & Chandler, 1988). We observed that removal of the vitelline envelope (VE) did not alter the SF-response (Fig 2B) and this is consistent with previous reports (Althaus et al, 2007). Next, oocytes without VE were treated with elastase (0.5 units/ml) or hyaluronidase (300 units/ml) to degrade elastin or hyaluronic acid, both ubiquitous ECM-components and involved in cellular mechanotransduction (Humphrey et al, 2014). After treatment with the enzymes, the cRNAs encoding α-, β-, and γENaC were injected and 24 hours later the oocytes were used for electrophysiological measurements and processed for electron microscopy (Suppl Fig 4). Due to the fragility of oocytes without VE and with an enzymatically impaired ECM, SF-responses were assessed by switching from low to high SF (0.01 to 0.2 instead of 0 to 0.2 dyn*cm^−2^), to prevent flow-induced ruptures of the oocytes.

**Figure 2.**
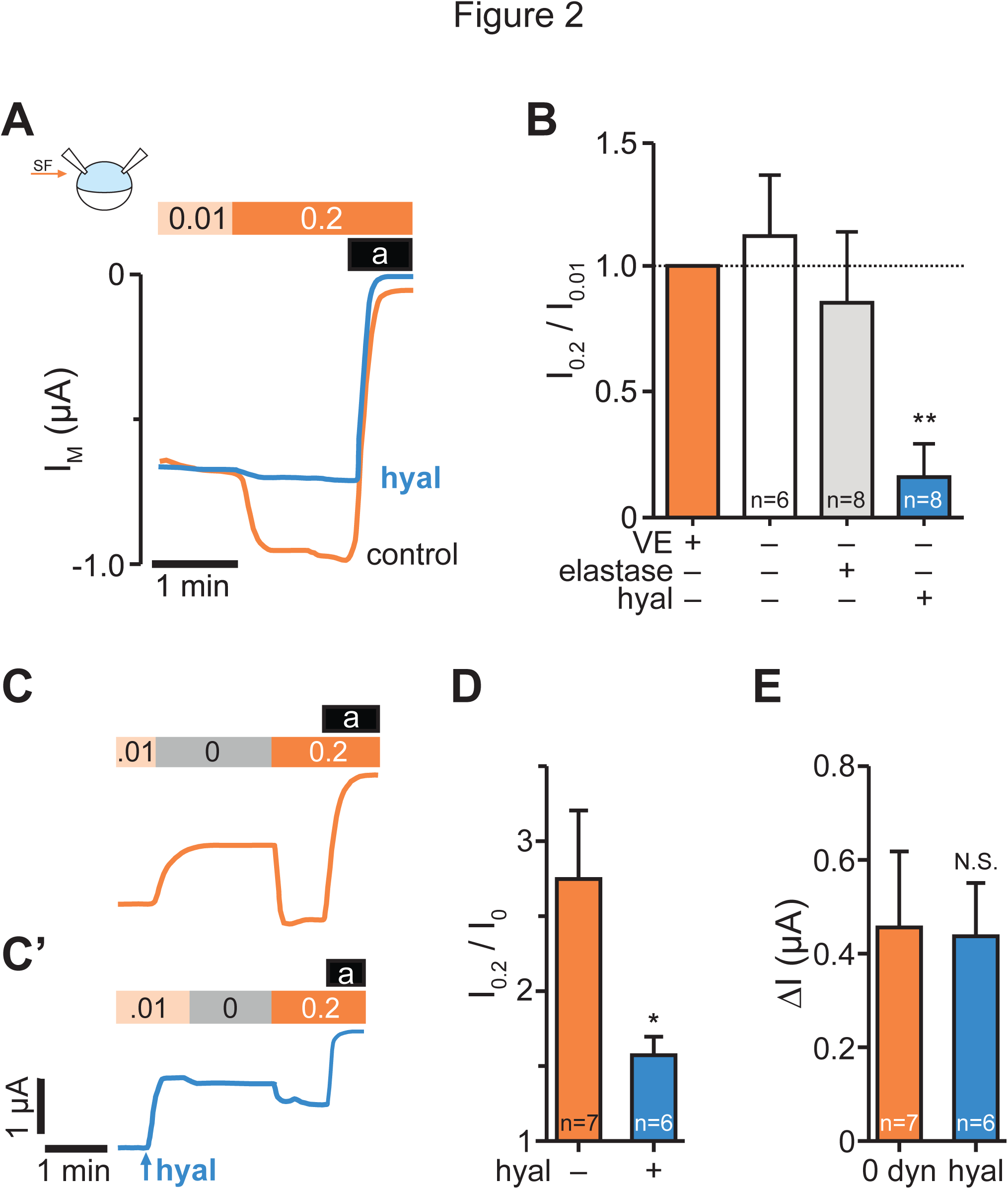
**Degradation of hyaluronic acid abolishes the SF-response of ENaC.** A SF responses observed in devitelinised ENaC-expressing oocytes that were either untreated (control, orange line) or additionally incubated with hyaluronidase (hyal, blue line). SF was applied by increasing the bath perfusion (0.01 → 0.2 dyn*cm^−2^). B Averaged SF-responses (I_0.2_/I_0.01_) from differentially treated oocytes to degrade certain ECM components (VE: vitelline envelope). SF-responses were normalized to corresponding control oocytes (intact ECM, orange bar) using cells from the same animals. The SF-response was largely decreased in hyaluronidase-treated oocytes (**: *P* < 0.01, one sample *t*-test, two tailed). C Stopping the bath perfusion (0.01 → 0 dyn*cm ^−2^) induced a current decrease and the following application of SF (0 → 0.2 dyn*cm^−2^) resulted in a current increase. C The acute application of hyaluronidase (300 units/ml) without changing SF (SF = 0.01 dyn*cm^−2^) decreased the current. Subsequent perfusion stop (0.01 → 0 dyn*cm^−2^) did not further affect the current. The subsequent application of SF (0 → 0.2 dyn*cm^−2^) resulted in a small SF-response. D SF-responses (I_0.2_/I_0_) from experiments shown in panel C and C’. Hyaluronidase decrease the SF response (*: *P* < 0.05, two tailed unpaired *t*-test). E Current changes induced by perfusion stop (0 dyn) and acute hyaluronidase application (hyal) had similar effects on the current (N. S., not significant, two tailed unpaired *t*-test).

Degradation of elastin did not alter the SF-response compared to non-treated oocytes (Fig 2B). By contrast, the treatment with hyaluronidase significantly reduced the SF-response (Fig 2A, B). It is unlikely that hyaluronidase impaired ENaC function directly since the cRNAs encoding for the ENaC-subunits were injected after the incubation with hyaluronidase. Also, the baseline amiloride-sensitive currents of hyaluronidase-treated oocytes (0.5 ± 0.06 µA, n = 8) were similar to those of untreated oocytes (0.6 ± 0.03 µA; n = 8; *P* = 0.8, two tailed unpaired *t-*test).

For further characterization, hyaluronidase (300 units/ml) was acutely applied to ENaC-expressing oocytes while measuring the I_M_. Hyaluronidase, applied through perfusion with 0.01 dyn*cm^−2^, decreased the I_M_ by approx. 47 ± 9 % within 20-30 seconds (Fig 2C’, E). Subsequent switch from 0.01 to 0 dyn did not have an effect (n = 7; *P* = 0.3 comparing values before and after 0 dyn, two tailed paired *t*-test) and there was a small effect in response to 0.2 dyn (Fig 2C’). In corresponding measurements without hyaluronidase, the switch from 0.01 to 0 dyn resulted in a decrease of I_M_ of 49 ± 7 % and the subsequent SF application resulted in an obvious effect (Fig 2C). From the experiments depicted in Fig 2C and 2C’ two things should be noted: (1) the change in current (∆I) observed with hyaluronidase was almost identical compared with eliminating SF (switch from 0.01 to 0 dyn; results summarized in Fig 2E). (2) The SF response was reduced by hyaluronidase (Fig 2D). These considerations are further supported by experiments showing that hyaluronidase had no effect when applied in the absence of SF (Suppl Fig 5). In summary, these experiments highlight the interdependent activity of hyaluronidase and SF for the mechanosensitive activity of expressed ENaC. They also confirm that the ECM is an essential component for mechanical regulation of ENaC.

### SF-mediated regulation of ENaC in arteries depends on the endothelial glycocalyx

To further confirm SF-induced activation of ENaC in a relevant mammalian physiological model, experiments with isolated pressurized and intraluminally perfused murine carotid arteries were performed. Here, low SF (flow rates < 100 µl/min) caused a slight reduction of the vessel diameter followed by an increase in response to elevated SF (Fig 3A) that is in agreement with flow-mediated vasodilation (Davies, 1995). Amiloride (10 µM) prevented the vasoconstriction with low SF and augmented the vasodilatory effect in response to elevated SF (Fig 3A). This indicates that SF-mediated regulation of ENaC contributes to regulation of the vascular tone. The potential contribution of the ECM for SF-sensing of ENaC in arteries was also addressed by experiments with hyaluronidase. In arteries, hyaluronic acid is an important part of the endothelial glycocalyx, a luminal ECM of endothelial cells (Reitsma et al, 2007). The glycocalyx is also crucial for the sensation of SF in blood vessels and the regulation of the vascular tone (Davies, 2009; Hahn & Schwartz, 2009).

**Figure 3.**
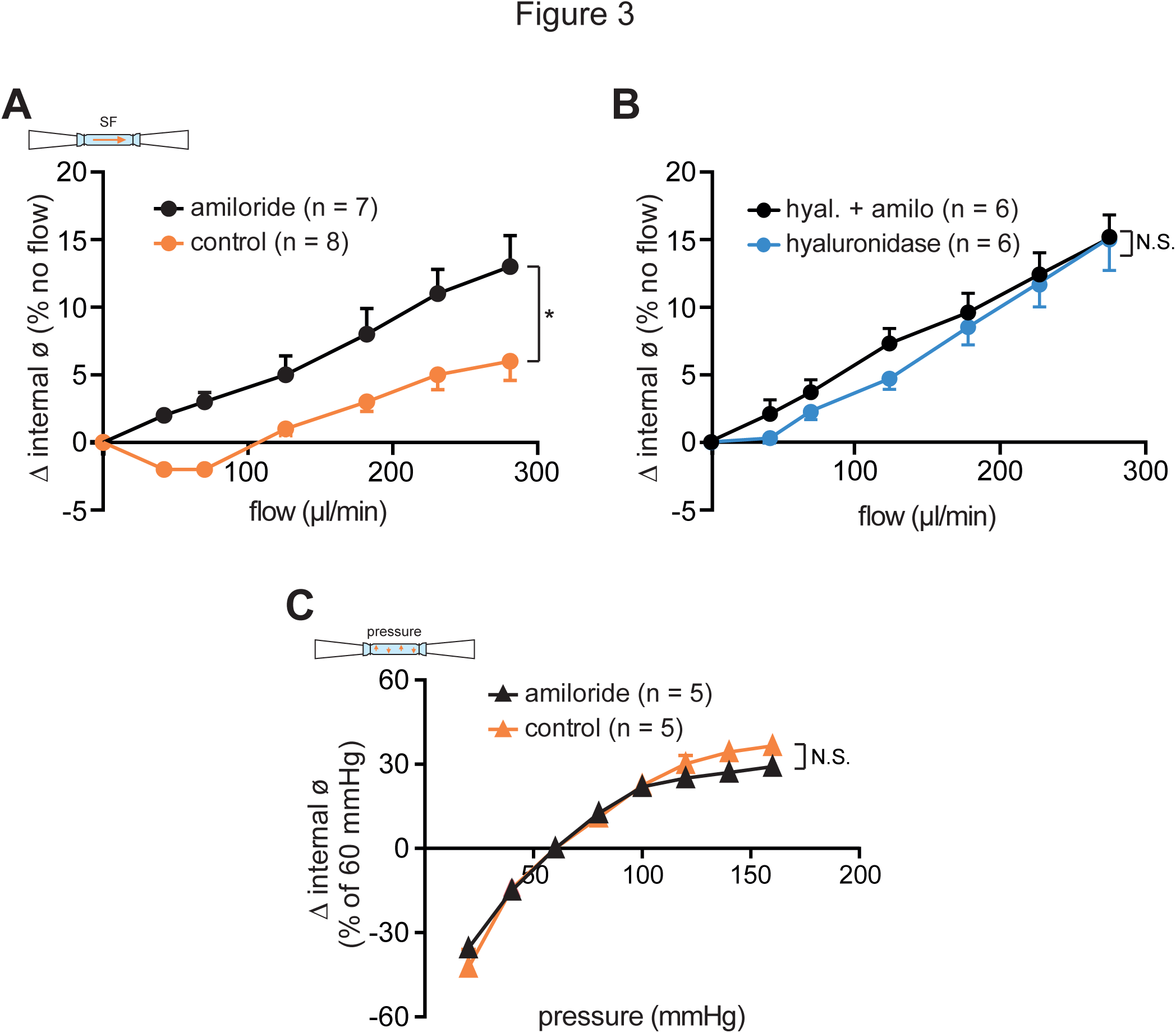
**ENaC-mediated regulation by SF in arteries depends on the endothelial glycocalyx.**. A Effect of SF on isolated pressurized carotid arteries from mice at mean arterial pressure of 60 mmHg. Increased intraluminal flow affected the internal diameter of the arteries under control conditions. Intraluminal perfusion of amiloride (10 µM) augmented the SF-induced increase of the vessel diameter. B Flow-induced changes after treatment with hyaluronidase to remove the endothelial glycocalyx. Note that amiloride had no effect, indicating an impaired ENaC activity in response to SF due to removal of the ECM. C Results from pressure myography experiments depicting changes of the internal diameter of carotid arteries at different intraluminal pressures. Amiloride had no effect indicating that ENaC does not mediate pressure responses in carotid arteries. Mean values ± SEM; *: *P* ≤ 0.05; N. S., not significant; repeated measures 2-way ANOVA)

Isolated arteries were treated with hyaluronidase and subsequently exposed to increased intraluminal flow, both in the absence and presence of amiloride (10 µM). The flow-induced vasodilation of hyaluronidase treated arteries was similar to the response observed with amiloride in non-hyaluronidase treated arteries and importantly, no response to amiloride was detected in hyaluronidase treated arteries (Fig 3B). This indicates an interdependent activity of ENaC and the ECM in response to SF in isolated arteries.

Additional experiments were performed to assess whether an amiloride-sensitive component could be observed in response to increased intraluminal pressure. No amiloride effect was detected in response to increased intraluminal pressure (Fig 3C). This indicates that ENaC in carotid arteries responds to SF rather than pressure. Further, these experiments confirm that SF-dependent activity of ENaC in arteries depends on an intact ECM/endothelial glycocalyx.

So far, regarding the first aspect of our study our experiments confirm for the first time that an intact ECM is required for mechanical activation of ENaC in response to SF – in an expression system as well as in freshly isolated arteries. These results also support the concept that an intact ECM is required for the transduction of force to regulate ENaC/DEG channel function.

### Extracellular *N*-glycans of asparagines are important for SF sensation

As it is common for all ENaC/degenerin proteins, the extracellular domains (EDs) of the ENaC subunits constitute approx. 70 % of the entire protein (Kashlan & Kleyman, 2011), and thus represent likely anchor points for a tether that can interact with the ECM.

Since the αENaC subunit is known to be crucial for the formation of fully functional channels (Canessa et al, 1994b) we focused on the ED of this subunit to identify a potential tether. Also, *N*-glycans of proteins were described to be important for cell-cell interactions and intercellular adhesion involving the ECM (Varki, 2007). Therefore, we hypothesized that *N*-glycosylated asparagines of αENaC could represent suitable anchor points and that the attached carbohydrates (representing the *N*-glycans) could be tethers that facilitate the interaction with the ECM for SF sensation and transduction.

Based on the consensus-sequence NXS/T (N: asparagine, X: any amino acid except proline, S: serine, T: threonine) for *N*-linked glycosylation (Spiro, 2002) and a sequence alignment with the known glycosylation sites of rat αENaC (Canessa et al, 1994a; Snyder et al, 1994) (Suppl Fig 6), five asparagines within the ED of the human αENaC subunit likely to be *N*-glycosylated were identified (N232, N293, N312, N397 and N511). Homology modelling of the human αENaC-subunit revealed that these asparagines are localized on the outer protein surface (Fig 4A, which is in accordance with previous findings from mouse ENaC (Kashlan et al, 2011)), and could provide access to the ECM. To challenge our hypothesis replacement of these asparagines should impair the SF-response of ENaC.

**Figure 4.**
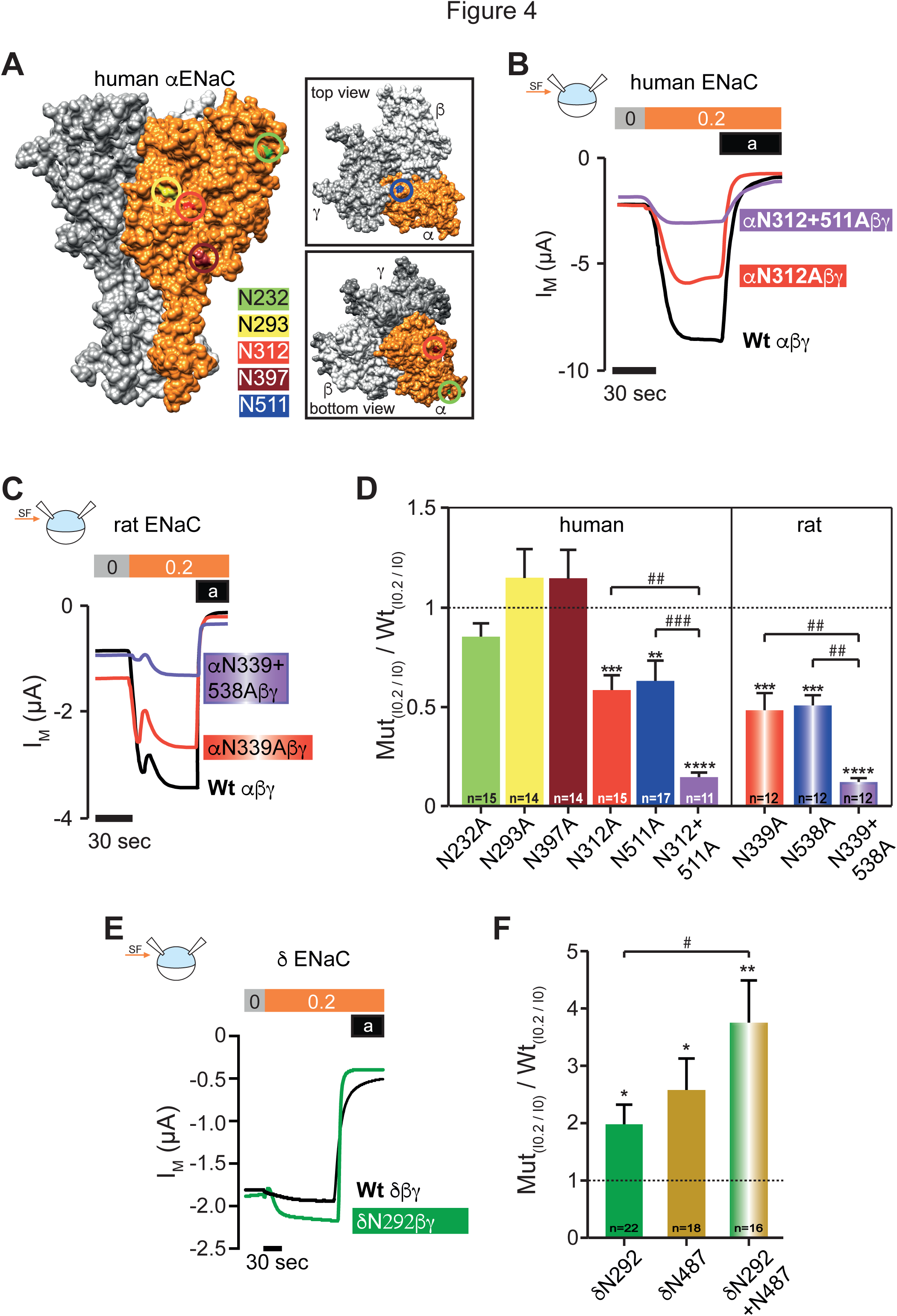
**Extracellular *N*-glycosylated asparagines are important for the SF response.** A Connolly surface model of a human αβγENaC-heterotrimer in the open state, displaying the predicted localization of glycosylated asparagines 232 (green), 293 (yellow), 312 (red), 397 (brown) and 511 (blue) within the α-subunit (highlighted in orange). These asparagines (N) were replaced by alanines (A) using site-directed mutagenesis. B Current-traces of the SF effects in oocytes that express either human wild type (Wt), a single (αN312A) or double mutation (αN312+511A) of the α subunit together with β and γ. C Replacement of the conserved corresponding asparagines of the rat αENaC subunit (N339 and N538) confirmed that these two asparagines are important for SF-sensing. D Replacement of certain asparagines of human and rat α ENaC decreased the SF response. Dashed line represents SF responses from corresponding control experiments using either human or rat Wt αβγENaC. E Experiments depicting the SF-responses of channels consisting the δ subunit. The insertion of a glycosylation motif increased the SF response (green line). F The SF effects obtained with δ ENaC subunits that contained additional *N*-glycosylation motifs were increased. The δ subunits were co-expressed with human β and γ ENaC. In D and F: Relative SF responses (I_0.2_/I_0_) of mutated channels were normalized with respect to Wt channels indicated by the dotted line (means ± SEM). \**P* < 0.05, \*\**P* < 0.01, \*\*\**P* < 0.001, \*\*\*\**P* < 0.0001 compared to Wt by one sample *t*-test, two tailed; #*P* < 0.05 ##*P* < 0.01 and ###*P* < 0.001 as indicated by one way ANOVA with Bonferroni´s multiple comparison test.

Each of the five asparagines was individually substituted with alanine and the modified αENaC proteins (αN232A, αN293A, αN312A, αN397A and αN511A) were co-expressed with wild-type (Wt) β and γ ENaC subunits. The responses to SF of these channels was determined and compared with those of Wt channels. No change of the SF effect was observed with αN232A, αN293A or αN397A (Fig 4D). In contrast, the SF-responses of channels consisting of αN312A or αN511A subunits were significantly reduced (Fig 4B, D). Simultaneous replacement of both asparagines further reduced the SF response (αN312+511Aβγ, Fig 4B, D), indicating that N312 and N511 contribute independently to SF-sensing. However, no additive effects were observed when N312 was mutated in combination with N232, N293 or N397 (Suppl Fig 7). Since asparagines 312 and 511 are conserved across mammalian orthologs we expected that replacement of these asparagines in the αENaC subunit of another species would provide similar results. For this approach, the corresponding asparagines of the rat αENaC ortholog (N339 and N538) were replaced with alanines, and these subunits were expressed with Wt rat β and γENaC. As observed with the human αENaC subunit, both asparagine mutations individually reduced the SF-response and simultaneous mutation had an additive effect (Fig 4C, D).

The role of *N*-glycans in mediating mechanical activation of ENaC was further investigated in experiments using the δENaC subunit (SCNN1D). The δENaC subunit can replace the α subunit to form a functional channel with β- and γENaC (Waldmann et al, 1995). SF activates δβγENaC, but the magnitude of the SF-response is smaller compared with αβγENaC (Abi-Antoun et al, 2011). A potential explanation for this could be the lack of *N*-glycosylated asparagines that correspond to N312 and N511 of αENaC. We decided to insert *N*-glycosylated asparagines into the δENaC. Locations were chosen that correspond to N312 and N511 of αENaC, and we hypothesized that this would produce δβγ channels that are more responsive to SF. To address the hypothesis, two modified human δENaC subunits were generated. A *N*-glycosylation motif (NNS) was inserted between amino acids 291 and 292 (δN292, corresponding to N312 of αENaC) and three amino acids (LPH) were replaced at position 487-489 by NYT (δN487, corresponding to position 511 of αENaC). The resulting channel subunits δN292 and δN487 were individually expressed with Wt human β and γENaC and exposed to SF. Both channels were activated more strongly by SF compared with the channel containing the Wt δ subunit. A third modified δ subunit carrying both glycosylation motifs responded even more strongly to SF compared with the single mutations (Fig 4E, F).

These results show that removal of existing *N*-glycosylated asparagines in the ED of αENaC decreases the SF-response, whereas the insertion of *N*-glycosylated asparagines in the ED of δENaC increases the SF responsiveness of ENaC. These observations highlight the role *N*-glycosylated asparagines for the transduction of mechanical force and suggest that the attached *N*-glycans are molecular components of tethers required for the interaction with the ECM.

### Glycosylation of N312 and N511 are required for SF-sensation but do not influence basic channel function

To confirm *N*-glycosylation of N312 and N511 of the human αENaC subunit, immunoblotting experiments with C-terminal hemagglutinin (HA)-tagged human αENaC-constructs (HA-Wt, HA-N312A, HA-N511 and HA-N312+511A) were performed. These experiments were accompanied by electrophysiology experiments to determine whether the HA-tag affected the response to SF, and we observed that the SF-responses were similar compared with the untagged channels (Suppl Fig 7). For immunoblotting protein extracts of oocytes expressing either Wt HA-αβγ, HA-αN312Aβγ, HA-αN511Aβγ or HA-αN312+511AβγENaC were lysed and proteins detected by anti-HA antibody. To observe deglycosylated Wt HA-αENaC protein, the samples were treated with peptide-*N*-glycosidase F (PNGase F). The glycosylated Wt HA-αENaC was detected with a relative molecular weight (M_R_) of about 80 kDa (Fig 5A) and this size is in accordance with previous findings (Harris et al, 2008). The deglycosylated Wt HA-αENaC subunit was observed at approximately 70 kDa (Fig 5A). HA-αENaC-mutants containing a substitution of a single asparagine displayed a reduced M_R_ (about 77 kDa, Fig 5A) compared with HA-Wt. Replacement of both asparagines further decreased M_R_ to about 74 kDa. This confirms N312 and N511 as glycosylation sites within the ED of human αENaC since removal of the asparagines and thus the *N*-glycans reduces the molecular mass of the ENaC subunits.

**Figure 5.**
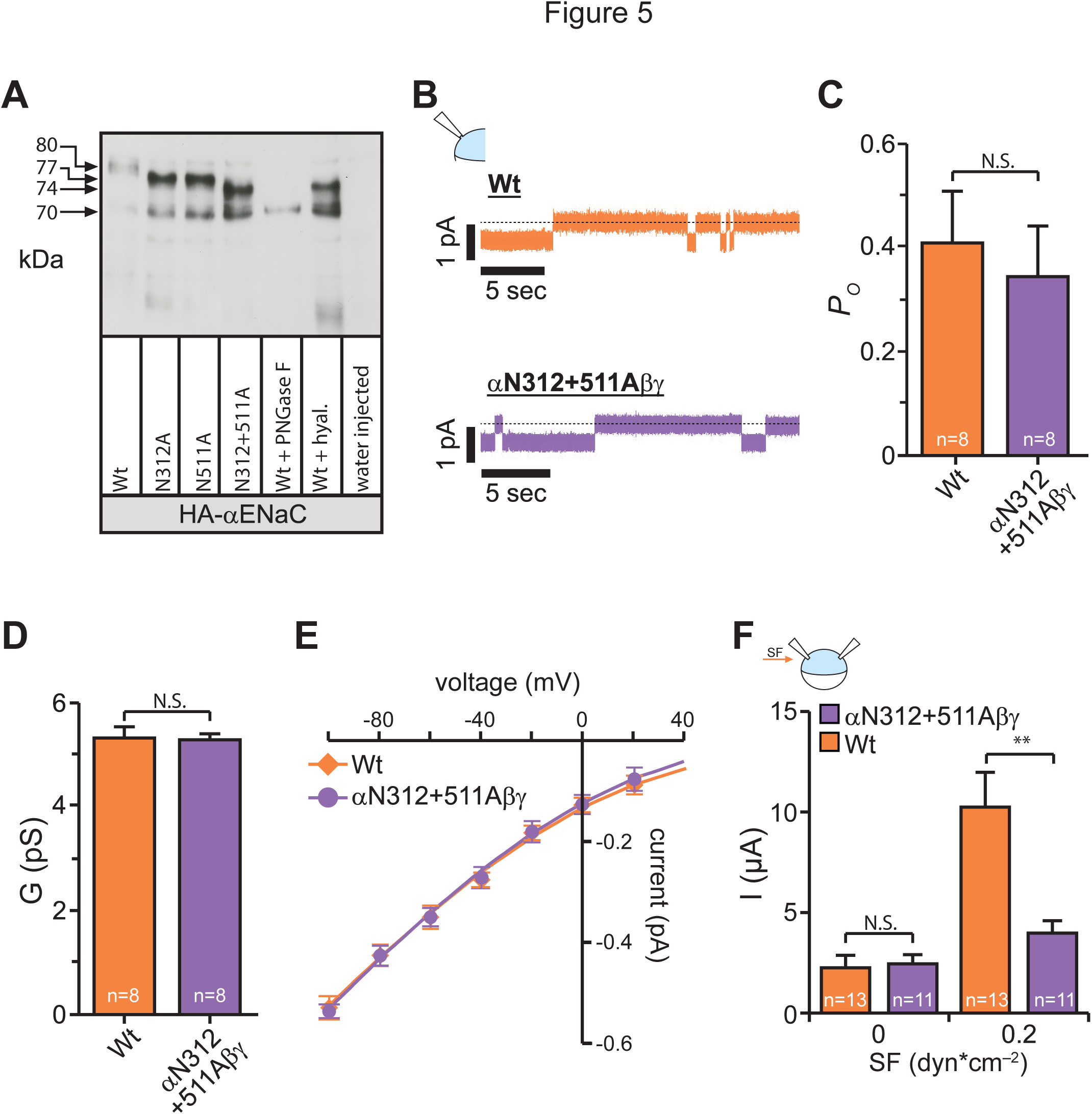
**Replacement of asparagines eliminates glycosylation and impairs the SF-response but does not alter basal channel function.** A Immunoblotting of whole-cell lysates from oocytes expressing either HA-tagged human Wt-αENaC (Wt), single-(αN312A or αN511A) or double *N*-mutants (αN312+511A), each in combination with the β and γ subunit. The shift in relative molecular mass (M_R_) is observed between the Wt (approx. 80 kDa) and *N*-mutants (approx. 77 kDa for N312A and N511A and approx. 74 kDa for N312+511A) corresponding to a lack of glycosylation. Removal of *N*-linked glycosylation by PNGase F resulted in the complete loss of the upper(glycosylated) band and shifted the band to approx. 70 kDa (Wt+PNGase F). Treatment of Wt with hyaluronidase resulted in a band of approx. 74 kDa (Wt+hyal.). Cell lysates of water-injected control-oocytes served as a control for antibody specificity. The depicted blot is representative for n = 6. B Current traces of cell attached single-channel recordings at a membrane-potential of −100 mV using devitellinised oocytes. C The open probability (P_O_) was not affected by replacement of the asparagines (N. S., not significant, two tailed unpaired *t*-test). D The single channel conductance (G) was also not affected by simultaneous mutation of N312 and N511 N.S., not significant, two tailed unpaired *t*-test). E The current-voltage relationship was not changed by the mutations. Values represent means ± SEM of at least nine current deflections for each voltage step obtained from at least three different recordings and cells. Means were fitted by the Goldman-Hodgkin-Katz equation. The calculated permeability for Na^+^ was identical (Wt: 1.72×10^−12^ cm^−3^ s^−1^; αN312+511AβγENaC: 1.73×10^−12^ cm^−3^ s^−1^). F TEVC experiments were performed to compare whole cell currents of Wt-and αN312+511AβγENaC. Without SF, ENaC-mediated currents (determined by amiloride Application) were similar between Wt-and αN312+511AβγENaC, implying a comparable number of active channels on the cell surface. However, the subsequent SF responses (0 → 0.2 dyn*cm^−2^) revealed a significantly decreased SF-response of αN312+511AβγENaC compared with the Wt. Data in C, D and F: Means ± SEM. N. S., not significant; ** *P* < 0.01; two tailed unpaired *t*-test.

To determine whether replacement of N312 and N511 influences basic channel function beside its impact on the SF response, single channel patch-clamp experiments were performed. Cell-attached recordings from oocytes expressing αN312+511Aβγ confirmed that neither the open probability, single channel conductance nor permeability for Na^+^ was different to the Wt-αβγ channel (Fig 5B-E). In addition, whole-cell amiloride-sensitive currents of Wt-αβγENaC and double-mutant αN312+511Aβγ were similar in the absence of SF (Wt: 2.3 ± 0.6 µA, αN312+511Aβγ: 2.4 ± 0.6 µA, *P* = 0.9; two tailed unpaired *t*-test, Fig 5F), whereas the SF-response was significantly reduced by replacement of N312 and N511. These observations confirm that neither membrane expression nor basic function of the channel is affected by the asparagine replacements supporting the exclusive role of these *N*-glycosylated asparagines for SF-sensation. It further supports the concept that the *N*-glycans are crucial for mechanical activation, potentially as components of tethers required for the connection to the ECM.

## Discussion

From the experiments performed in this study we clearly obtained evidence that an intact ECM is required for mechanical activation of ENaC since enzymatic degradation of hyaluronic acid impaired the response to SF in both *Xenopus* oocytes and isolated intraluminal perfused carotid arteries. These findings extend our knowledge about the role of the ECM for mechanotransduction of ENaC, and identify, for the first time, the contribution of glycosaminoglycans of the ECM for mechanical activation of ENaC/DEG channels.

However, our results do not prove that ENaC is directly attached to hyaluronic acid for two reasons: (1) hyaluronidase cleaves chondroitin and chondroitin sulfate in addition to hyaluronic acid (Menzel & Farr, 1998) and (2) hyaluronic acid might serve as a ‘backbone’ to which other ECM-components such as proteoglycans are attached (Kiani et al, 2002). Therefore, degrading hyaluronic acid may impair the integrity of the entire ECM, which could secondarily affect the integrity of another ECM component that may serves as ENaC’s direct anchor point. Although it remains unknown which ECM component is connected to ENaC, our experiments highlight the role of glycosaminoglycans for mechanotransduction and provide further support for the ‘force-from filament’ principle (Katta et al, 2015), where the ECM serves as the extracellular filamentous ‘sensing’ structure for mechanosensitive ion channels.

The EDs of ENaC and DEG proteins comprise a large portion of the protein and they are known to be important for regulating channel activity (e.g. ENaC activity through proteolytic cleavage (Kleyman et al, 2009)). Our study may provide further explanations for the size of the EDs as they may also reach out so far to enable a connection with the ECM. *N*-glycans attached on the top of the ED can further increase the distention by another 6 nm (Rudd et al, 1997) supporting the idea of the ED (anchor point) and their *N*-glycans (tethers/part of tether) as components for mechanotransduction.

We have evidence that *N*-glycans of two asparagines within the ED of the αENaC subunit are involved in SF-sensing. The *N*-glycans may facilitate the connection between the sensing structure and the channel as the ‘receiver’. Replacement of certain asparagines of αENaC, and the subsequent loss of *N*-glycans, impaired the SF-mediated activation of rat and human ENaC. Moreover the introduction of *N*-glycosylation motifs into δENaC increased the channel’s response to SF. This provides strong support for the concept that *N*-glycans could be crucial components of tethers that facilitate a physical connection between the channel and other components of the ECM. This is in agreement with the role of *N*-glycans of integrins (Janik et al, 2010; Gu et al, 2012), which are transmembrane proteins involved in mechanotransduction by facilitating interactions with the ECM (Schwartz, 2010). The proposed new role of *N*-glycans as facilitators of mechanical activation of ion channels is also supported by studies identifying that glycan/glycan interactions provide a high binding strength that facilitates binding of bacteria to host cells (Day et al, 2015), but also cell adhesion in multicellular organisms (Popescu et al, 2003).

Based on the resolved structure of acid sensing ion-channel 1 (ASIC1) – another member of the ENaC/DEG protein family that is considered to be mechanosensitive (Chen & Wong, 2013) – it is believed that the structure of the ED of αENaC resembles an outstretched hand (subdivided into thumb, palm, finger and knuckle domains) encircling a ball (β-ball,(Jasti et al, 2007)). The connection between the ED and the transmembrane segments – and thus the gating machinery – is provided by the wrist-domain, the pre-M2 segments and the β-turn (Kashlan et al, 2011; Shi et al, 2013). Recently, different domains of ENaC subunits including the thumb (Shi et al, 2011), finger (Shi et al, 2012a), second transmembrane segments (Abi-Antoun et al, 2011), pore-region (Carattino et al, 2004; 2005), pre-M2 segments (Carattino et al, 2005), wrist (Shi et al, 2012b) and β-turn (Shi et al, 2011) were shown to be involved in SF-dependent activation of ENaC. Whether these domains are directly involved in SF sensation or convey SF-induced conformational changes (downstream of SF-sensing) to alter channel gating remains unknown. However, homology modelling of the αENaC subunit revealed that N312 is localized on the outer surface of the palm-domain (Fig 6A) and N511 on top of the knuckle-domain (Fig 6B). Neither, the palm-nor the knuckle-domain were previously investigated with regard to SF activation.

**Figure 6.**
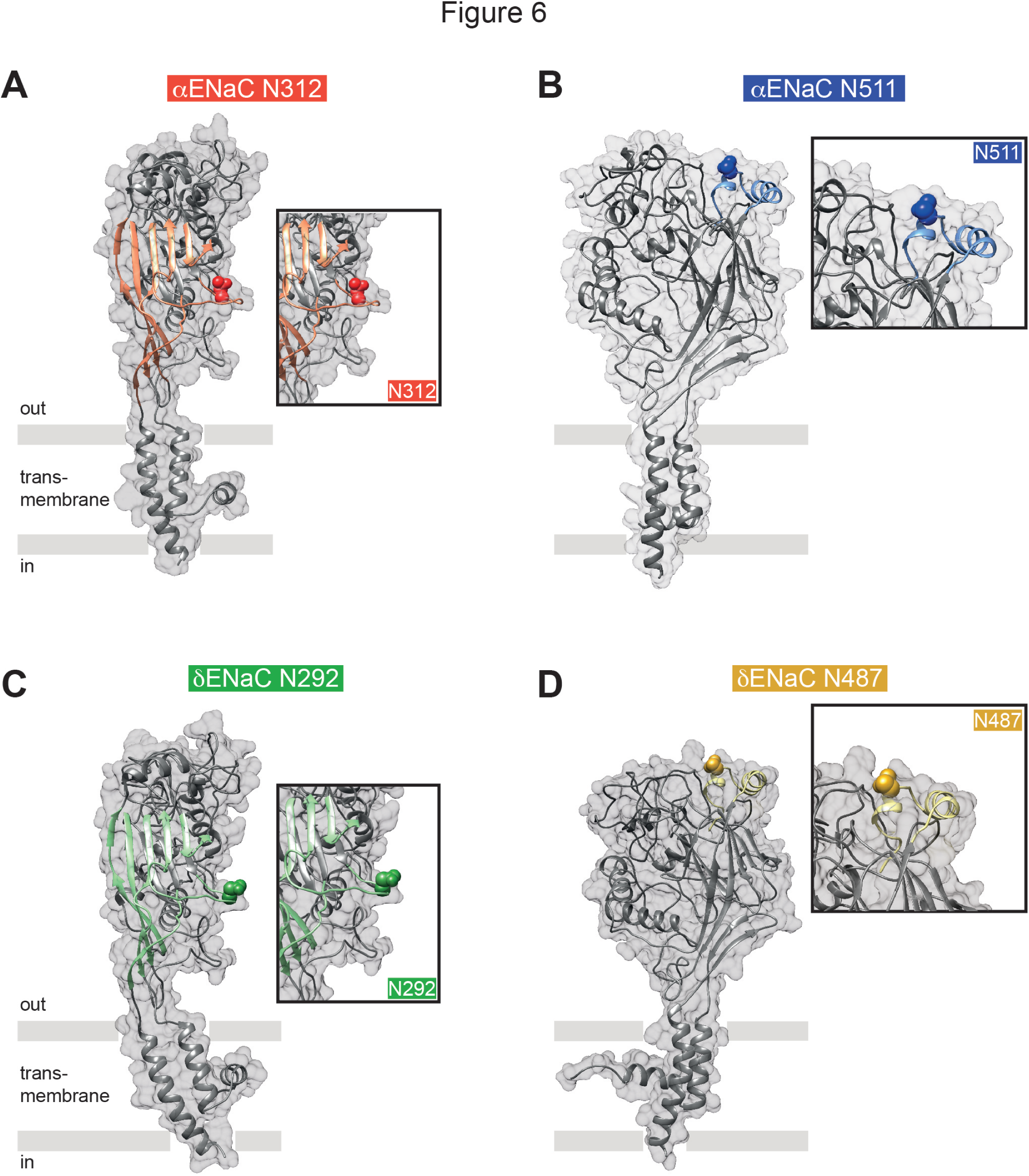
**The asparagines responsible for the SF effect are located within the palm and knuckle domain**. A Model of the human αENaC-subunit, highlighting asparagine 312 (red) within the palm domain (accentuated in coral). B Localization of asparagine 511 (blue) on top of the knuckle domain (sky blue). C Model of the human δENaC-subunit containing the inserted glycosylation site at position 292 (green). D The δENaC subunit with the glycosylation site at position 487 (olive). The positions of the asparagines were predicted by homology modelling based on the resolved ASIC1 structure in the open state. Our model indicates that the glycosylated asparagine residues are localized at exposed positions, likely to provide connections to the ECM.

### Physiological role of interdependent ENaC/ECM activity

The results observed in our study could be of considerable relevance for various physiological processes and blood pressure regulation in particular. Hyaluronic acid is a major component of the endothelial glycocalyx (Reitsma et al, 2007) comprising the luminal ECM of endothelial cells and is crucial for mechanotransduction of SF in blood vessels (Davies, 2009; Hahn & Schwartz, 2009). Here SF regulates the production and release of the vasodilator nitric oxide (NO) (Davies, 1995). This mechanotransduction mechanism enables local autonomous modulation of vascular tone by adjusting the vessel diameter to alterations in blood perfusion rates (Davies, 1995). Strikingly, vascular ENaC exerts its effects by influencing NO production (Oberleithner et al, 2007; Perez et al, 2009). Therefore, our study and the observations in carotid arteries in particular provide the first evidence for a functional interaction between the endothelial glycocalyx as the force sensing structure, and endothelial ENaC as the receiver molecule that transduces mechanical SF into a cellular signal. This underlines the importance of mechanotransduction beyond the perception of touch and pain.

### Concluding remarks

To our knowledge, this study is the first to deliver an explanation for the, as yet unknown, interaction between a mechanosensitive ion channel with the ECM or other components. Our data provide unique evidence to support the ‘force-from filament’ principle for ENaC/DEG that depends on channel *N*-glycans (Fig 7). This discovery may also apply to other ion channels such as ASICs and ATP-gated P2X4 channels that share a similar structure (Jasti et al, 2007; Gonzales et al, 2009; Kawate et al, 2009) and are suggested to form mechanosensitive channels (Yamamoto et al, 2006; Chalfie, 2009; Kessler et al, 2011). Further, it may be proposed that *N*-glycans are uniform structures for mechanotransduction that may also apply to channels with a different structure like PIEZO1 (Ge et al, 2015) and TRP channels. PIEZO1 is an interesting candidate since it has been identified as shear sensor in the vasculature (Li et al, 2014), as well as a stretch activated channel that is associated with the cytoskeleton (Cox et al, 2016). Interestingly, three peripheral extracellular domains within PIEZO1 are required for mechanical activation and this trimeric structure is reminiscent of trimeric channels (Akyuz & Holt, 2016; Zhao et al, 2016), that is common for members of the ENaC/DEG protein family (Jasti et al, 2007; Chen et al, 2015). It may be hypothesized that a trimeric structure equipped with *N*-glycans is a unifying structural feature for mechanotransduction of ion channels.

**Figure 7.**
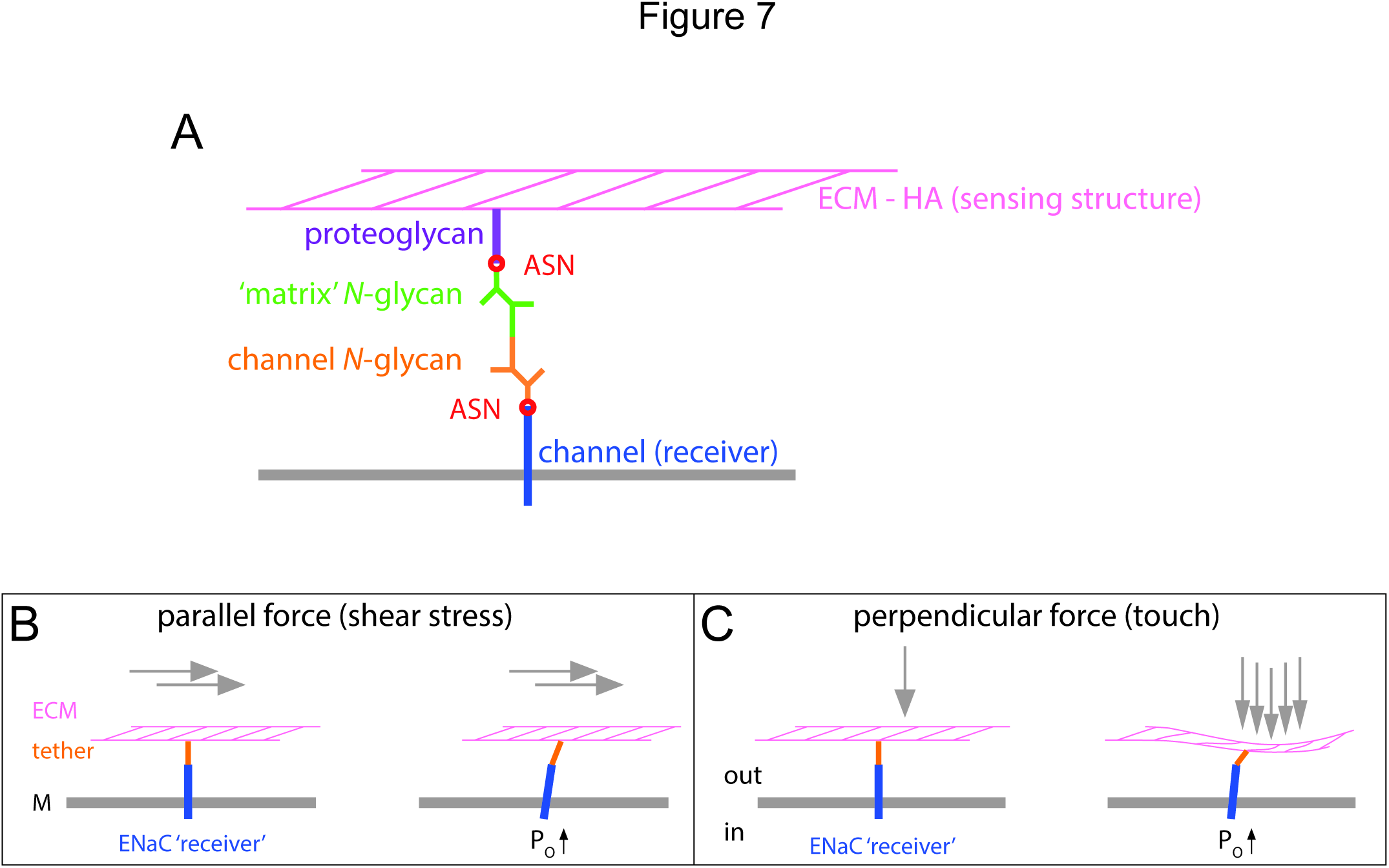
**Force transduction through *N*-glycan ‘tethers’ and the ECM.** A Proposed structure and architecture of an extracellular tether based on *N*-glycan/*N*-glycan interactions. B Shear force caused by fluids flowing over the surface of cells can cause deflection of the ECM. Through a tether involving N-glycan interactions this can induce a conformational change of the channel to increase its activity. C The application of a force perpendicular to the ECM/cell surface can also induce shear movement affecting the tethered channel in a similar way. ECM: extracellular matrix; HA: hyaluronic acid; ASN: *N*-glycosylated asparagine; M: membrane; P_O_: open probability.

It may also be considered that the role of extracellular glycans is not limited to the sensation of shear force caused by the flow of liquids over cell surfaces. *N*-glycan mediated activation of channels can also facilitate the sensation of touch (Fig 7). Deflection of the ECM in response to touch can cause a relative movement of the matrix with respect to the membrane. This relative movement of the two layers can produce shear force on a structure that is attached to both layers. This would activate the same mechanism as observed in response to laminar flow. It would also explain the activation of ENaC in renal arteries subjected to elevated pressure. Here ENaC proteins of vascular smooth muscle cells mediate a vasoconstriction in response to extension of the vessel walls (Jernigan & Drummond, 2005; 2006). Deformation of the vessel wall may result in shear movement of the matrix in between adjacent smooth muscle cells.

Based on our results we propose that *N*-glycosylated asparagines and particularly the attached *N*-glycans, are a new unifying structural feature of mechanotransduction. This can have far-reaching impact on our understanding of how mechanical forces regulate ion channels activity in conjunction with the ECM.

## Materials and Methods

### Heterologous expression of human ENaC in *Xenopus* oocytes

Human ENaCs were heterologously expressed in *Xenopus laevis* oocytes. The oocytes, harvested from adult female South African clawed frogs (*Xenopus laevis*) were defolliculated enzymatically by incubation (90 minutes) in collagenase-containing (1.5 mg/ml) CulORi-solution (CulORi: cultivation oocyte Ringer’s solution; composition in mM: 90 NaCl, 1 KCl, 2 CaCl_2_, 5 HEPES (4-(2-hydroxyethyl)-1-piperazineethanesulfonic acid), 2.5 Na^+^-pyruvate, 0.06 penicillin, 0.02 streptomycin, pH 7.4) and subsequent incubation (10 min) in a Ca^2+^-free oocyte Ringer (ORi; composition in mM: 90 NaCl, 5 HEPES, 1 KCl, 1 EGTA (ethylene glycol tetraacetic acid), pH 7.4). Oocytes of stages V and VI were sorted and subsequently used for cRNA-injection. cRNA encoding the human or rat α, β and γ ENaC-subunits were dissolved in DEPC (diethylpyrocarbonate)-treated water and injected at a ratio of 1:1:1 (0.9 ng human and 2.1 ng rat) RNA/oocyte) *via* a micro injector (3-00-203-XV, Nanoject, Drummond Scientific, Broomall/USA). Control oocytes were injected with corresponding volumes of DEPC-treated water. After injection, oocytes were cultured in a low-Na^+^-solution (in mM: 10 NaCl, 80 NMDG (*N*-methyl-*D*-glucamine), 1 KCl, 2 CaCl_2_, 5 HEPES, 2.5 Na^+^-pyruvate, 0.06 penicillin, 0.02 streptomycin; pH 7.4) at 16 °C. The culture-solution was changed every day. Two-electrode voltage-clamp (TEVC)-recordings were performed at room temperature within 24 – 48h after injection.

### Two-electrode voltage-clamp recordings (TEVC)

Oocytes were placed in a customized flow chamber, which was connected to a pressure-controlled perfusion system (ALA Scientific Instruments, Westbury, NY/USA). The flow-chamber had a narrow channel for perfusion, designed to allow a consistent application of SF to the oocyte’s surface and a rapid solution exchange. The oocytes were positioned in front of the inflow (2 mm drilled hole). SF was applied by increasing the perfusion velocity *via* elevating the air pressure on the perfusate. This allowed for a precise application of SF-rates in physiologically relevant ranges (0.001 to 1.3 dyn*cm^−2^).

Chlorinated silver wires served as intracellular microelectrodes and were mounted in pulled borosilicate glass capillaries (Cat NR 1103245, Hilgenberg, Malsfeld/Germany), filled with 1 mM KCl. The membrane potential of the oocytes was clamped to −60 mV using a TEVC-amplifier (Turbo-TEC03X or Turbo-TEC05, NPI, Tamm/Germany). The transmembrane current (I_M_) was recorded *via* a strip chart recorder (model BD 111, Kipp&Zonen, Delft/The Netherlands) and monitored by a voltmeter (model DMT 7976, düwi, Mömbris/Germany). The entry of Na^+^-ions is defined as negative inward current. An increase of an inward current corresponds to a downward deflection of the measured I_M_. For determining the ENaC-mediated current, amiloride (10 µM) was applied at the end of each experiment. The difference between I_M_ and the current that remained in the presence of amiloride (I) represents the amiloride-sensitive, ENaC-mediated current (I_ami_ = I − I_M_) and was used for statistical analysis.

### Site-directed mutagenesis

Plasmid constructs (in pTNT, Promega, Mannheim/Germany) containing the coding sequence for human α, δ or rat α ENaC subunit (with and without C-terminal HA-tag) were used as templates for site-directed mutagenesis. Mutations were introduced using the Quikchange Lightning site-directed mutagenesis kit (Agilent Technologies, Waldbronn/Germany) according to the manufacturer’s instructions. Primer sequences for the mutations are listed in Table 1 (human) and Table 2 (rat). Double mutants were generated sequentially. Mutated plasmids were transformed into competent DH5α *E. coli* cells and isolated by miniprep (Perfectprep Miniprep Kit, 5PRIME, Hilden/Germany). Successful mutagenesis was confirmed by DNA sequencing (Eurofins, Ebersberg/Germany). cRNA was generated from the mutated plasmids by *in vitro* transcription (RiboMAX Large Scale RNA-production system-T7, Promega) according to the manufacturer’s instructions and then used for oocyte-injection.

**Table 1:**
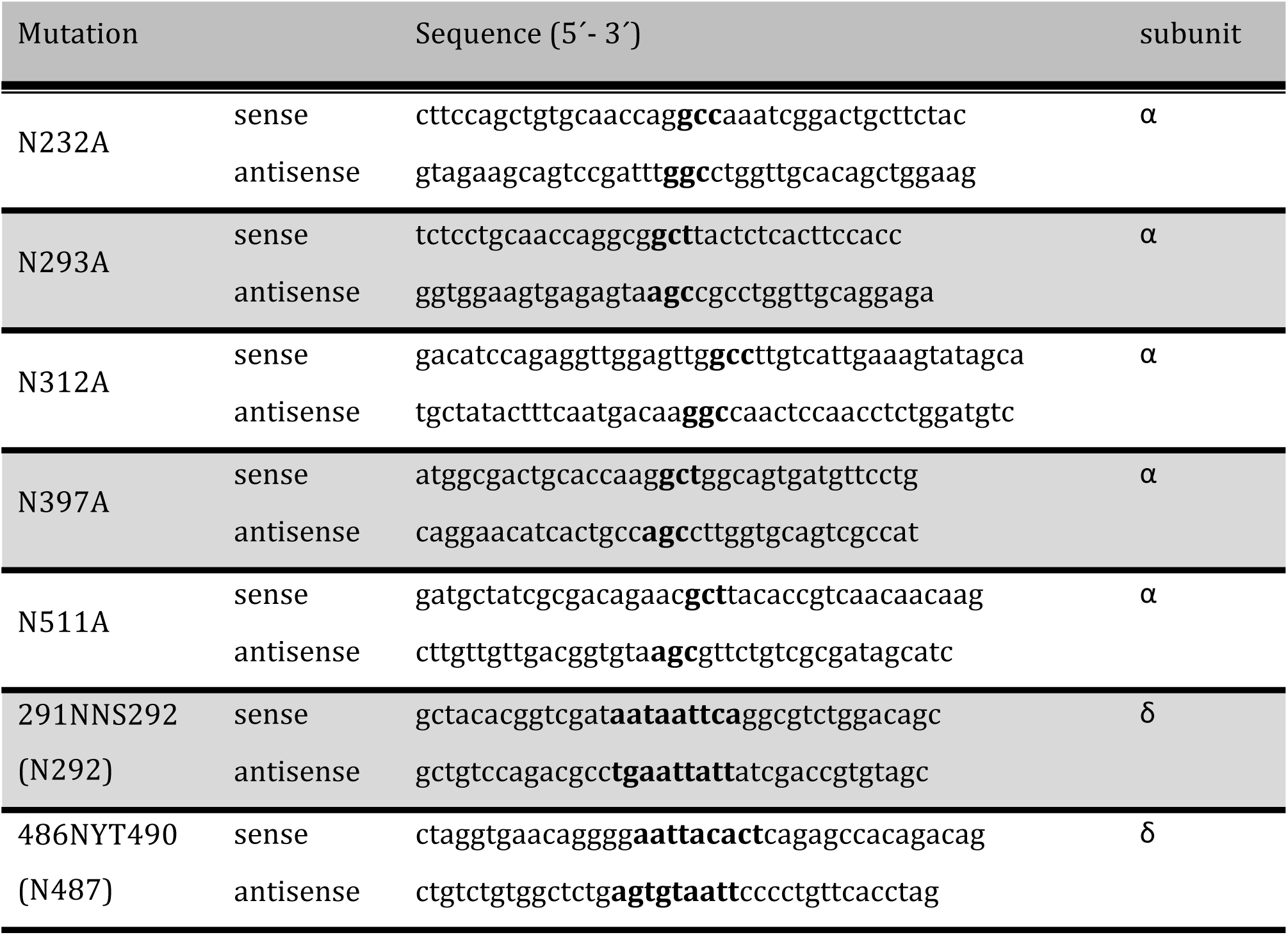
Primer sequences used for human ENaC site-directed mutagenesis.

Mutated nucleotides to replace asparagines (αENaC) and to insert *N*-glycosylation motifs (δENaC) are bold and underlined.

**Table 2:**
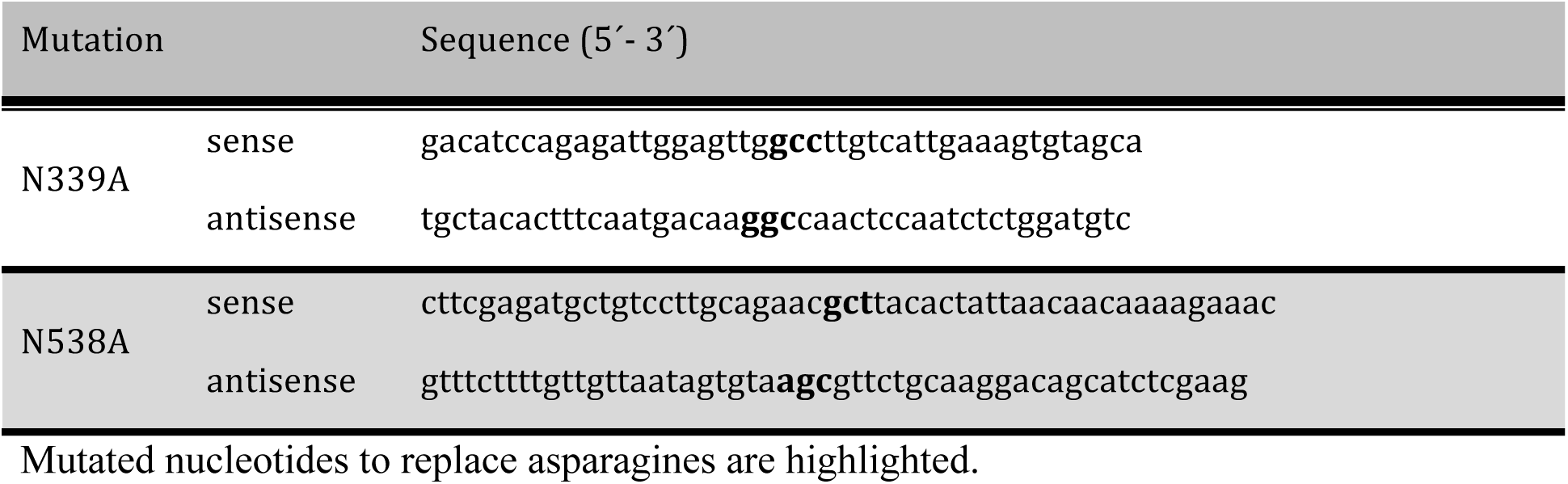
Primer sequences used for rat αENaC site-directed mutagenesis

Mutated nucleotides to replace asparagines are highlighted.

### Proposed structures of human α and δENaC by homology modeling

The chicken ASIC1 crystal structure in the open state (PDB: 4NTX, (73)) was chosen as the template for creating a predicted 3D model of the human α and δ ENaC subunit as well as the heterotrimeric αβγENaC. Therefore, amino acid sequences of human α (UniProt identifier: P37088-1), β (UniProt ID: P51168-1), γENaC (UniProt ID: P51170-1) were used as input and aligned to ASIC1 (UniProt ID: P78348) sequence using Clustal Ω with the BLOSUM90 scoring matrix. 15 different models for each subunit were generated (Modeller 9.16, http://www.salilab.org/modeller/). The model with the lowest normalized Discrete Optimized Protein Energy (zDOPE)-score was selected as the best model and used for further structure analysis. For δENaC the respective *N*-glycosylation motifs were inserted manually into the δENaC sequence (UniProt ID: P51172-1) at the appropriate positions and subsequently used as input for homology modeling. Predicted structures of each ENaC subunit were aligned to the structure of the appropriate subunit of the ASIC1 homotrimer (PDB: 4NTX.pdb2.gz) using UCSF Chimera 1.10.2 (http://www.cgl.ucsf.edu/chimera; Resource for Biocomputing, Visualization, and Informatics, University of California, San Francisco, supported by NIGMS P41-GM103311 (74)). To indicate the putative location of selected asparagines, the ‘surface’ function was used to visualize the solvent-accessible surface area (Connolly surface).

### Determination of ENaC glycosylation by western blot

The glycosylation-states of Wt-αENaC and *N*-mutants were characterized by western blot. Protein extracts were obtained from oocytes expressing C-terminal HA-tagged αENaC (Wt or mutant) in combination with β and γ ENaC subunits. Oocytes were homogenized in lysis buffer (10 µl per oocyte; in mM: 83 NaCl, 10 HEPES, 1 MgCl_2_, 1 % Triton-X 100; pH 7.9, supplemented with EDTA-free protease inhibitor cocktail; Roche, Mannheim/Germany). Lysates were cleared by centrifugation for 10 min at 1000 g and 4 °C. The aqueous protein-containing phase was transferred into a new tube using a 2 ml syringe in combination with a cannula tip. Lysates were centrifuged again and remaining yolk was removed in the same manner. Protein concentrations were determined with Bradford-reagent (Applichem, Darmstadt/Germany) and 30 µg protein was generally utilized for experiments.

In order to remove *N*-linked glycosylation, Wt ENaC protein extracts (30 µg) were treated with PNGase F (New England Biolabs, Frankfurt a. M./Germany) according to the manufacturer’s protocol. Protein extracts were combined with loading dye (Rotiload 1, Roth, Karlsruhe/Germany) and loaded on 8 % polyacrylamide gels. For deglycosylation with hyaluronidase, Wt-ENaC-extracts (30 µg) were supplemented with loading dye, heated to 85 °C for 10 min and subsequently supplemented with hyaluronidase (final concentration 300 units/ml). Samples were incubated for 1 hour at 30 °C. All other protein extracts were combined with loading dye and heated to 85 °C for 10 min before being separated by SDS-PAGE (sodium dodecyl sulfate-polyacrylamide-gel-electrophoresis). Proteins were transferred onto nitrocellulose-membranes (Bio-RAD, Munich/Germany) and blocked for 1 hour with 5 % BSA (bovine serum albumin) in TBST (Tris buffered solution with 0.1 % Tween 20; in mM: 150.6 NaCl, 15.2 Tris-HCl, 4.6 Tris-Base, pH 7.6). αENaC was detected with antibodies directed against its C-terminal HA-tag (1 mg/ml; dilution 1:10.000; HA Epitope Tag Antibody, Thermo Scientific, Waltham, MA/USA) by incubating the membrane over night at 4 °C. After washing 3 times in TBST, membrane was incubated with a peroxidase conjugated secondary antibody (0.6 mg/ml; dilution 1:3000; Pierce Rabbit Anti-Mouse IgG, (H+L), Thermo Scientific Waltham, MA/USA) for 1 hour at room temperature. Bands were visualized using a customized ECL (enhanced chemiluminescence)-solution and captured on photo reactive films (Amersham Hyperfilm ECL, GE Healthcare, Pittsburgh, PA/USA).

### Removal of extracellular matrix components of *Xenopus* oocytes

The extracellular matrix of *Xenopus* oocytes consists of 1) the vitelline envelope (VE) and 2) filamentous, glycocalyx-like structures within the perivitelline space (PS) (Larabell & Chandler, 1988). To remove the VE, oocytes were incubated in CulORi-solution, which was supplemented with mannitol (10 mg/ml). Due to the hyperosmolarity of this solution, oocytes shrank within 5 minutes, whereas the surrounding VE retained its structure. This enabled the mechanical removal of the VE using two fine forceps. Oocytes were subsequently transferred into the recording chamber and TEVC recordings were performed in accordance with previous studies (Althaus et al, 2007) (Vitzthum et al, 2015).

ECM-components within the PS were degraded enzymatically. To remove the oocyte hyaluronic acid network, devitellinised oocytes were incubated for 2 hours at 30 °C in a CulORi-solution (pH 6) supplemented with 300 units/ml hyaluronidase (ROTH, Karlsruhe/Germany). For elastin-degradation, devitellinised oocytes were incubated at 37 °C for 30 min in CulORi-solution (pH 8) supplemented with 0.5 units/ml elastase (Sigma Aldrich, Seelze/Germany).

## Electron microscopy

### Transmission electron microscopy (TEM) of *Xenopus* oocytes

After TEVC-measurements, *Xenopus* oocytes were fixed in 0.15 M HEPES-buffer (Calbiochem, Darmstadt/Germany) supplemented with 1.5 % glutaraldehyde (Plano GmbH, Wetzlar/Germany) and 1.5 % paraformaldehyde (Merck, Darmstadt/Germany). Afterwards, oocytes were washed (2 x 10 minutes) in 0.15 M HEPES-buffer, postfixed for 1 hour with 1 % osmium tetroxide (Merck, Darmstadt/Germany) before being washed in distilled water (2 x 10 minutes). Subsequently, the oocytes were block-fixed in half-saturated uranyl acetate-solution (Merck, Darmstadt/Germany) overnight and washed in distilled water (3 x 10 minutes). Dehydration of oocytes was achieved by incubation in an ascending ethanol chain (30, 50, 70, 90 and 100 %, Merck, Darmstadt/Germany). Afterwards, they were laid in a mixture of propylene oxide (Merck, Darmstadt/Germany) and epon (ratio 1:1, Plano, Wetzlar/Germany), finally embedded in pure epon and dried for 48 hours at 60 °C. Semithin (0.75 µm) and ultrathin (0.075 µm) sections were cut using a diamond knife and a microtome (Leica, Bensheim/Germany), placed on copper nets and contrasted with uranyl-acetate and lead citrate (Leica Reichert Ultrastainer, Bensheim/Germany). The ECM of the oocytes was examined using a transmission electron microscope (Zeiss EM 902, Oberkochen/Germany).

### Slam-freezing and TEM of HEK293 cells

HEK293 cells were cultured in growth medium (DMEM buffer with 4.5 g/l D-glucose, L-glutamine, 110 mg/l sodium pyruvate, 1 % MEM non-essential amino acids, 1 % penicillin/streptomycin) with 10 % fetal bovine serum in a 5 % CO_2_ atmosphere at 37 °C. Rapid specimen freezing was performed using a Reichert-Jung KF80 Plunge Freezer device with metal mirror (MM80) attachment (Leica Microsystems, Vienna/Austria). Frozen cells were transferred to a EM AFS2 automatic freeze-substitution device (Leica Microsystems, Vienna/Austria) where they were freeze-substituted in acetone containing 1 % osmium tetroxide at –90 °C for 3 days and then the temperature was slowly raised to room temperature over 2 to 3 days. The cells were rinsed in acetone and embedded in epoxy resin. The samples were then cut perpendicular to the growth plane into 80 nm sections using a Reichert-Jung Ultracut E ultramicrotome (C. Reichert, Vienna/Austria). Sections were examined using a CM-100 transmission electron microscope (Philips, Amsterdam/Netherlands).

## Patch-clamp experiments

### Single-channel recordings

Single-channel recordings were performed in the cell-attached configuration on devitellinised ENaC-expressing oocytes. Pulled and fire-polished borosilicate glass capillaries (Cat NR 1100303, Hilgenberg, Malsfeld/Germany) with an outer diameter of 1.6 mm and resistances between 4 – 10 MΩ served as patch-pipettes and were filled with extracellular analogous solution (in mM 90 NaCl, 1 KCl, 2 CaCl_2_, 5 HEPES, pH 7.4). The bath solution contained in mM: 145 KCl, 1.8 CaCl_2_, 2 MgCl_2_, 5.5 glucose, 10 HEPES, pH 7.2 in order to keep the endogenous membrane potential (V_M_) close to 0 mV.

Single-channel currents were amplified using an LM/PC amplifier (List Electronics, Darmstadt-Eberstadt/Germany) and filtered at 100 Hz with a low-pass filter (Frequency Devices, Inc., Ottawa, IL/USA). Data were acquired at 2 kHz using an Axon interface (1200er series) in combination with the Axon Clampex software 8.0.3 (Axon Instruments, Union City, CA/USA). Data were analyzed with OriginPro9.0G (Originlab, Northhampton, MA/USA). Single-channel events were recorded for approx. 5 minutes at a membrane potential of −100 mV. Na^+^-conductance, open probability, and permeability were estimated as previously described (Althaus et al, 2007).

### Whole cell recordings on HEK293 cells

HEK293 cells were grown under the same culture conditions as used for electron microscopy imaging. At approx. 70 % confluency, cells were passaged and used for transient transfection (jetPRIME polyplus transfection reagent) in accordance with the supplier’s protocol. Cells were grown on poly-L-lysine-coated glass coverslips (13 mm diameter) placed in 24-well plates (cell density approx. 50.000 cells/well) and treated with the DNA mix (150 ng of plasmids encoding the α, β and γ human ENaC subunit, including 50 ng plasmid encoding enhanced green fluorescent protein). The supernatant was removed after 2-4 hours and the growth medium was supplemented with 10 µM amiloride. Experiments were performed within 18-32 hours after transfection. Only fluorescence positive cells that expressed an amiloride-sensitive current were used for experiment. Coverslips were either changed after an experiment, or after 45 minutes of usage. Whole-cell current was amplified with an Axopatch 200B amplifier and digitized with a Digidata 1440A (Molecular Devices, Sunnyvale/USA). The bath solution contained (in mM): 140 NaCl, 3 KCl, 2 CaCl_2_, 1 MgCl_2_, 10 D-glucose, 20 HEPES (pH 7.4). Electrodes were pulled from borosilicate capillaries (1.5 mm diameter) with resistances between 1.5 and 3 MΩ and filled with intracellular solution containing (in mM): 90 K-gluconate, 10 NaCl, 10 KCl, 1 MgCl_2_, 10 EGTA, 60 HEPES (pH 7.3). Amiloride (10 µM) was applied to inhibit ENaC-mediated currents. Experiments were performed using BT-1-TB perfusion chamber in combination with a cFlow V2.x 8-Channel Switch/Flow Control System (Cell MicroControls, Norfolk/USA). This system allowed accurate control and changes of perfusion rates for SF-application. Whole-cell currents were recorded at –60 mV at room temperature.

### Pressure myography

All procedures were approved by the University of Otago Animal Ethics Committee and conducted in accordance with the New Zealand Animal Welfare Act. Male mice (C57BL/6, 8-10 weeks old) were terminally anaesthetized with sodium pentobarbitone (150 mg/ml) and exsanguinated by removal of the heart. The carotid arteries were identified and their *in vivo* lengths measured. One artery was removed, placed in ice-cold buffer and mounted in the pressure myograph system (DMT, Aarhus/Denmark). Vessels were super-perfused at 1 ml/min with warmed Krebs-Henseleit solution of the following composition (in mM): 119 NaCl, 4.7 KCl, 25 NaHCO_3_, 1.2 KH_2_PO_4_, 1.2 MgSO_4_, 11.1 glucose, 0.03 EDTA, 1.6 CaCl_2_, pre-oxygenated with 5 % CO_2_ in 95 % O_2_. The same solution was used for intraluminal perfusion and pressurization. After mounting, the vessel was allowed to equilibrate (approx. 1 hour including pressurization of the vessel to 60 mmHg intraluminal pressure), followed by a standard ‘wake-up’ protocol to ensure viability of the vessel (noradrenalin, acetylcholine, KCl). Intraluminal SF was applied by changing the inlet and outlet pressure without changing the mean intraluminal pressure. The internal and external diameter of the vessels was captured by a camera (The Imagingsource, Bremen/Germany) attached to a microscope (Nikon, Tokyo/Japan) and the edge detection routine of the DMT software. The flow and resulting SF-rates were determined by a flow meter (DMT 162FM) integrated into the system. Amiloride (10 µM) and hyaluronidase (300 U/ml) were added into the perfusate. Hyaluronidase was incubated for 10 minutes intraluminally (pressurized at 60 mmHg, no flow) and removed after incubation.

### Statistical analysis

Data are presented as mean ± standard error of the mean (SEM). Numbers of individual experiments are declared as *n.* The number of animals from which oocytes were used was ≥ 3. Statistical analysis was performed with Prism GraphPad (La Jolla, California. USA). Electrophysiological data were analysed for Gaussian normal distribution using the Kolmogorov Smirnov test. Dependent values were compared by the paired Student’s *t*-test. Independent values that were normally distributed were compared by an unpaired *t*-test. Non-normally distributed values were compared with the Mann-Whitney *u*-test. Normalized values were compared with one sample *t*-test. For multiple comparisons, one-way ANOVA followed by appropriate post hoc tests was used. Statistical analysis of blood vessel diameters was performed by repeated measures 2-way ANOVA. In all cases a *P*-value of at least < 0.05 was considered as statistically significant.

## Acknowledgements

The authors would like to thank Gerd Magdowski (Institute of Anatomy and Cell Biology, Justus-Liebig-University, Giessen, Germany) for assistance with electron microscopy on oocytes; Allan Mitchell and Sharon Lequeux (Otago Centre for Electron Microscopy, University of Otago, Dunedin, New Zealand) for assistance with the slam freezing/cryosubstitution and imaging of HEK293 cells; and Siegfried Kristek (Institute of Animal Physiology, Justus-Liebig-University, Giessen, Germany), Leo van Rens and James Woods (Emtech, University of Otago) for manufacturing the flow chamber and the pressurized perfusion system. The valuable comments and suggestions provided by Drs Fiona McDonald, Kirk L. Hamilton and Boris Martinac during editing and finalizing of the manuscript are greatly acknowledged.

## Author contributions

The study was designed by F.K. and M.F.F.K. performed recordings with αβγ ENaC in Xenopus oocytes, did immune blotting, provided the channel structure and electron microscopy images of oocytes. Z.A. performed pressure myograph experiments on isolated arteries.

D.B. performed the experiments with δβγ ENaC and measured whole-cell currents of HEK293 cells. M.K. contributed electron microscopy images from HEK293 cells. P.P.S and M.A assisted with the homology modeling.

D.A.R. generated and provided the HA-tagged α ENaC construct. The manuscript was drafted and written by F.K. and M.F. P.P.S, M.A.D.A.R. and W.G.C provided substantial contribution during editing and finalizing of the manuscript.

